# Early microglial priming in Alzheimer’s disease revealed by ME-seq

**DOI:** 10.64898/2026.01.30.702946

**Authors:** Bohan Zhu, Anurupa Ghosh, Zhe Wang, Zhicong Liao, Michael Appiah, Jiayi Li, Mimi Zhang, Federico Di Tullio, Agata Kurowski, John Fullard, Elvin Wagenblast, Martin J. Walsh, Alison Goate, Panos Roussos, Ka Lung Cheung, Nan Yang, Sai Ma

## Abstract

Epigenetic modifications, particularly DNA methylation, change dynamically with aging and are implicated in Alzheimer’s Disease (AD), yet how methylation interfaces with transcriptional and chromatin regulation at single-cell resolution remains poorly understood. Progress has been limited by a lack of scalable technologies capable of jointly profiling these regulatory layers. Here, we present ME-seq, a highly scalable technologies capable of simultaneously profiling DNA methylation, gene expression, and chromatin accessibility, while achieving a 100-fold reduction in cost. We generated over 400,000 single-nucleus trimodal profiles from the aging and AD mouse brain across ages, producing the first such atlas of neurodegeneration. We found AD progression triggers pronounced, disease-specific shifts in cellular composition, characterized by accelerated epigenetic aging and the expansion of disease-associated microglia (DAM). Integrative analyses, including aging clocks, revealed that DNA methylation acts as an early priming layer preceding transcriptional activation with IRF1 identified as a methylation-sensitive transcription factor serving as a gatekeeper for DAM activation. Our results establish ME-seq as a transformative tool for large-scale epigenomic dissection, revealing DNA methylation as a primary coordinator of cell-state transitions in the aging brain.

**Graphic abstract:** 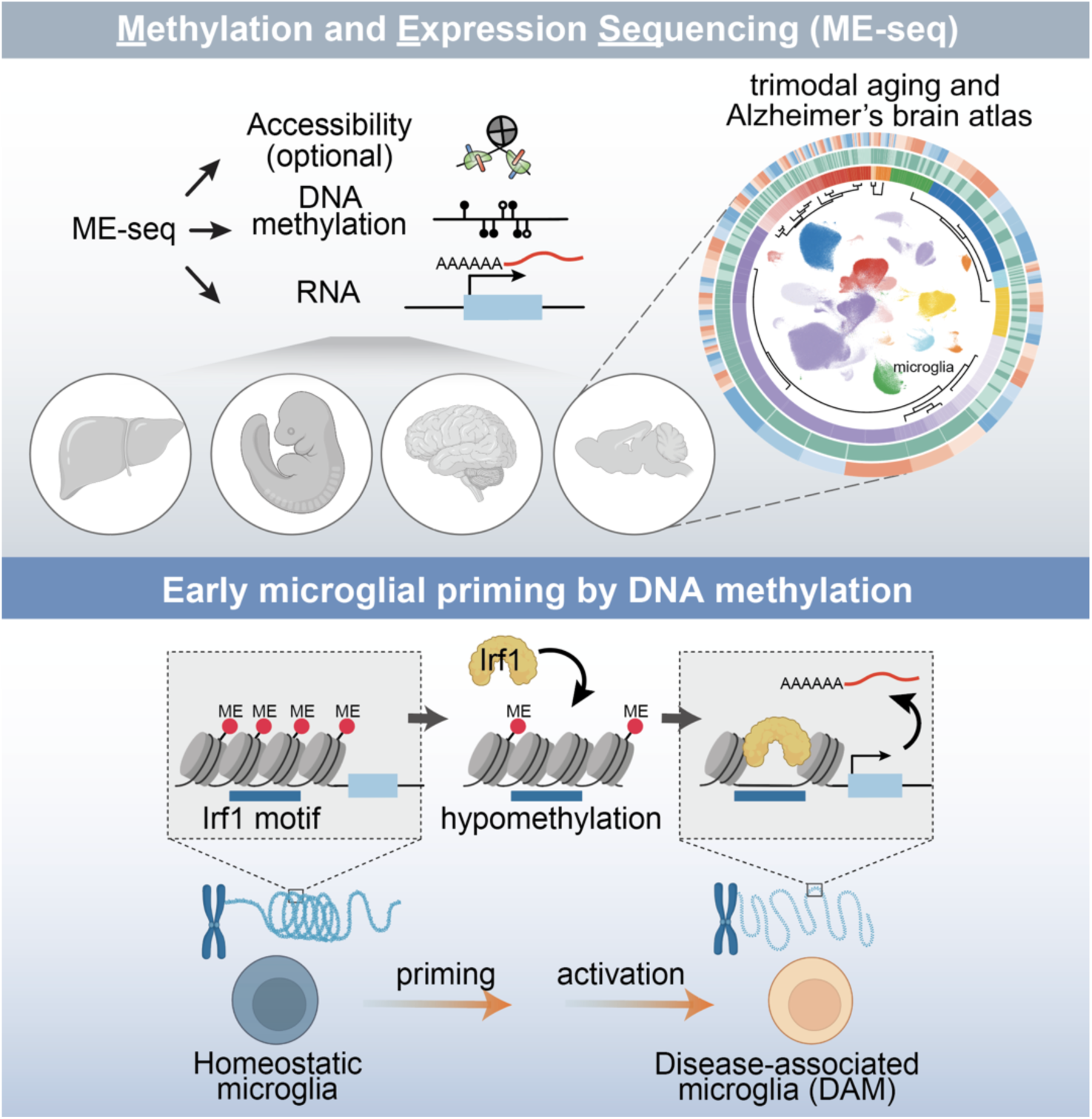

## Introduction

Over the past decade, a major scientific breakthrough has revealed that epigenetic modifications, particularly DNA methylation, evolve as we age^1^, serving as a molecular recorder of aging and a primary driver of neurodegenerative pathogenesis^2,3^ and cancers^4^. While aging remains the strongest risk factor for neurodegenerative diseases such as Alzheimer’s disease (AD) and Parkinson’s disease (PD)^5^, the logic that governs how homeostatic brain cells bypass epigenetic checkpoints to enter pathological states remains unresolved. Emerging studies suggest that brain aging is not only defined by cellular loss, but by a coordinated reprogramming of the DNA methylome and chromatin landscape, pushing astrocytes, oligodendrocytes, and microglia^6,7^ toward “primed” inflammatory states. However, the precise temporal sequence of these events—and whether epigenetic remodeling is a driver or a passenger of transcriptional dysregulation—remains a fundamental unanswered question in neurobiology.

AD is a global health crisis currently lacking effective disease-modifying therapies^8^. Mounting evidence implicates epigenetic dysregulation on DNA methylation^9–11^ and chromatin accessibility^12^ in AD pathogenesis, yet our understanding of the dysregulation is fundamentally limited by a resolution gap. Bulk-tissue studies provide an averaged signal that obscures the high-stakes regulatory shifts occurring in rare, transitional cell populations. While recent single-cell studies^13–15^ have begun to map gene expression and chromatin accessibility in the AD brain, they provide only a snapshot of active cellular states. They remain blind to the latent regulatory potential of the genome: the “priming” events that may occur when the methylome is remodeled before a gene is ever transcribed.

A major barrier to resolving this temporal hierarchy has been the lack of scalable technologies capable of joint multi-omic profiling. Current single-cell DNA methylation methods are cost-prohibitive and technically arduous, typically restricted to a few thousand cells per study^16,17^. This throughput is orders of magnitude below what is required to capture the continuum of microglial activation or to detect the subtle, cell-type-specific “epigenetic drift” that precedes neurodegeneration^18^. Additional complexity arises because DNA methylation signatures often do not align perfectly with known marker genes identified by RNA-seq, complicating data interpretation without paired transcriptomic data^19^. Furthermore, computational attempts to integrate independent “omic” datasets suffer from alignment artifacts, precluding the precise, cell-by-cell coupling of the methylome to the transcriptome needed to infer causality^20^.

This limitation is particularly consequential for the study of epigenetic priming, where DNA methylation changes may establish “transcriptional competence” during a silent lag phase before overt disease symptoms emerge. To resolve dynamics at scale, here we develop ME-seq, a high-throughput single-nucleus platform for the joint profiling of DNA methylation and gene expression, with the capacity for concurrent chromatin accessibility measurement (trimodal). By reducing the cost of single-cell methylome profiling by more than 100-fold, ME-seq enables the generation of atlas-scale datasets with the statistical power necessary to resolve rare regulatory transitions. We apply ME-seq to construct a trimodal single-cell atlas of the aging and AD mouse brain, encompassing over 400,000 single-nucleus profiles. Our analyses reveal that DNA methylation remodeling acts as a foundational priming layer, pre-configuring the transcriptional trajectories of disease-associated microglia (DAM) long before inflammatory activation occurs. Leveraging newly developed computational frameworks and identifying methylation-sensitive transcription factor (TF) networks and thousands of previously hidden *cis*-regulatory elements (CREs), we provide a mechanistically grounded framework for understanding the epigenetic origins of AD and establish ME-seq as a transformative tool for large-scale epigenetic discovery. These findings establish DNA methylation as a central driver of cell-state transitions in the aging brain and demonstrate the power of scalable, multimodal single-cell epigenomics to uncover regulatory mechanisms that are inaccessible to single-modality or low-throughput approaches.

## Results

### ME-seq enables massive-scale co-profiling of DNA methylation and gene expression

Understanding how DNA methylation interfaces with transcriptional regulation at single-cell resolution has been fundamentally limited by the “throughput-cost” barrier. While current multi-omic assays offer conceptual proof-of-concept, their scalability remains orders of magnitude lower than that of single-modality approaches, constraining their application to complex tissues or large-scale disease cohorts. To overcome these limitations, we developed ME-seq (**M**ethylation and **E**xpression co-assay with **seq**uencing), a highly scalable single-cell platform that jointly measures DNA methylation, gene expression, and chromatin accessibility within the same cell at an unprecedented scale.

ME-seq achieves this scalability by integrating TET-mediated enzymatic DNA methylation conversion^21^ (**Extended Data Fig. 1a**) with the combinatorial indexing framework of SHARE-seq^22^, enabling simultaneous, high-throughput capture of epigenomic and transcriptomic information. The platform is uniquely flexible, offering two complementary implementations: Open-ME-seq, which selectively interrogates DNA methylation within accessible regulatory elements for cost-effective large-scale mapping, and Closed-ME-seq, which utilizes lithium diiodosalicylate^23^ to disrupt nucleosomes, enabling unbiased whole-genome methylation profiling across even compacted chromatin regions (**Fig. 1a**).

**Figure 1:**
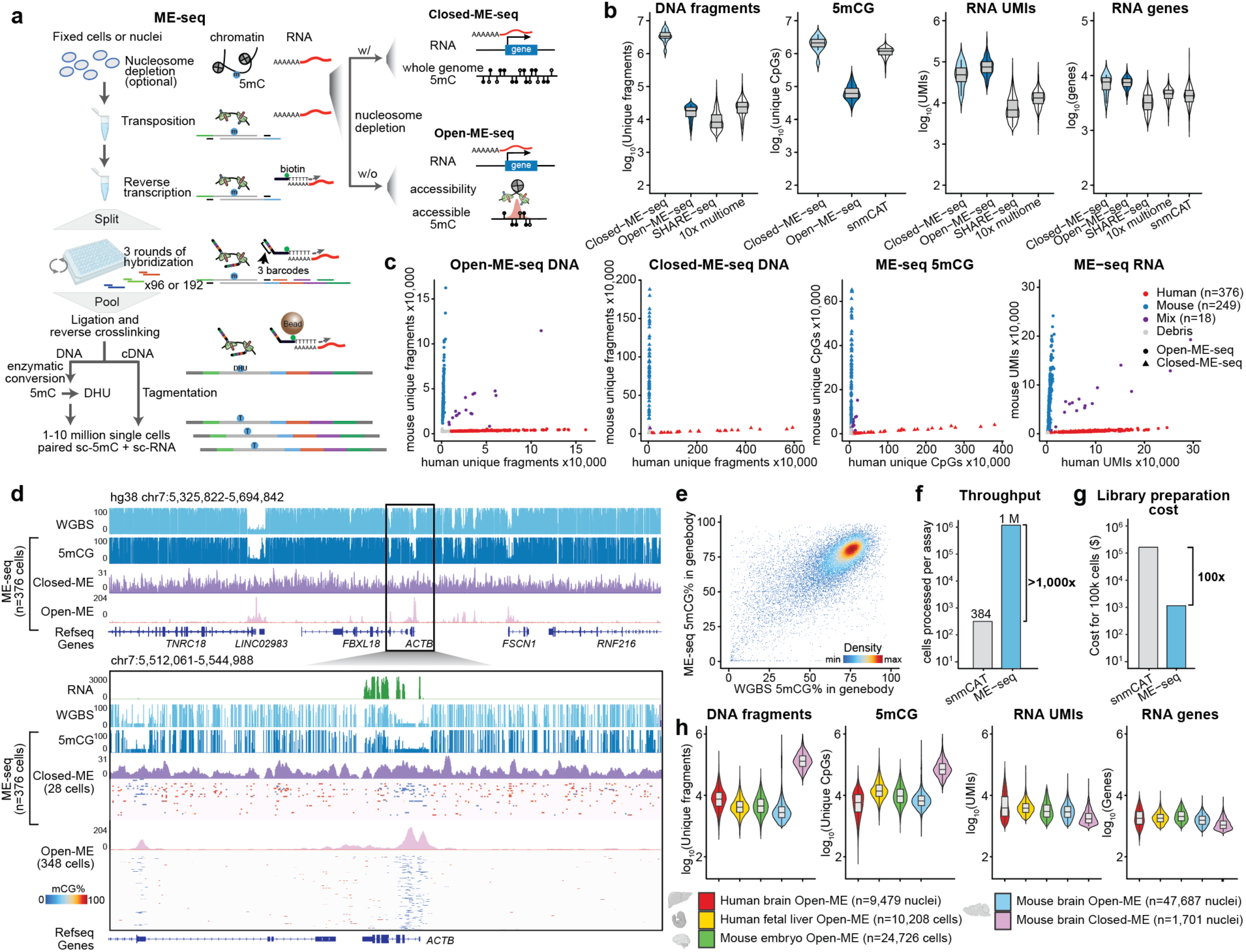
ME-seq enables massive-scale co-profiling of DNA methylation and gene expression. (a) Schematic diagram of ME-seq workflow. (b) Violin plots showing per-cell data yield for ME-seq compared with state-of-the-art single-cell multi-omic platforms (10x Genomics Multiome GSE178707 and snmCAT GSE140493) using HEK293 cells. Metrics include DNA fragments, covered CpG sites (5mCG), RNA unique molecular identifiers (UMIs), and detected genes per cell; only cells passing platform-specific quality control are shown. (c) Cross-contamination analysis using mixed human HEK293 and mouse NIH/3T3 cells. Each point represents one cell. From left to right: DNA libraries generated by Open-ME-seq, DNA libraries generated by Closed-ME-seq, DNA methylation profiles from ME-seq, and RNA libraries from ME-seq. Species-specific alignment quantifies human- and mouse-mapped reads to assess sample purity across assays. (d) Genome tracks around the housekeeping gene *ACTB*. Top, four tracks display: public HEK293 whole-genome bisulfite sequencing (WGBS) data; aggregated 376 ME-seq single cells; aggregated coverage from 28 Closed-ME-seq cells; and aggregated coverage from 348 Open-ME-seq cells. Bottom, zoomed views with additional single-cell DNA methylation level from 28 Closed-ME-seq single cells and 348 Open-ME-seq cells. (e) Comparison of gene body DNA methylation levels in public WGBS data and ME-seq, with each dot representing one gene. (f) Comparison of assay throughput between ME-seq and snmCAT, measured as the number of single-cell libraries generated per experiment. (g) Comparison of per-sample library preparation costs for ME-seq and snmCAT under the conditions used in this study, excluding sequencing costs. (h) Comparison of per-cell data quality across human brain, human fetal liver, and mouse embryo samples profiled with Open-ME-seq, together with mouse brain samples generated using both Open- and Closed-ME-seq, including unique DNA fragments, covered CpG sites, RNA UMIs, and detected genes per cell.

The architecture of ME-seq is engineered to preserve data integrity across modalities. Following fixation and Tn5 transposition of cells or nuclei, we perform *in situ* mRNA capture and reverse transcription using a dual-priming strategy (biotinylated poly(T) and random hexamers). Nuclei undergo three rounds of split-pool barcoding, generating up to 10^7^ unique combinations to ensure minimal collision rates during massive multiplexing (**Extended Data Fig. 1b, Supplementary Table 1**). A critical innovation is the post-barcoding decoupling step: cDNA is isolated via streptavidin affinity, while genomic DNA undergoes non-destructive enzymatic oxidation and pyridine borane reduction. Unlike traditional bisulfite treatment, this enzymatic strategy prevents DNA degradation, preserves library complexity^21^, and ensures that the rigorous conditions required for methylation conversion do not compromise RNA quality.

We performed extensive optimizations to ensure ME-seq meets the stringent data-quality requirements of the community (**Extended Data Fig. 1d-j**). First, we re-engineered the library structure for compatibility with the latest high-throughput Illumina Nova X Plus sequencing platform, reducing the sequencing cost by over 4-fold (**Extended Data Fig. 1a**). Second, we achieved a 95.0% 5mC conversion efficiency with a negligible 1.4% false-positive rate, matching the performance of gold-standard whole-genome bisulfite sequencing (WGBS) (**Extended Data Fig. 1d**). Third, we optimized nucleosome disruption and Tn5 transposition steps to maximize library complexity and signal-to-noise ratio (**Extended Data Fig. 1e, f**). Notably, the TSS enrichment (open chromatin bias) was eliminated upon lithium diiodosalicylate treatment, suggesting even genomic coverage (**Extended Data Fig. 1g**). Finally, by enriching for short cDNA fragments and optimizing reverse transcription, we enhanced RNA library complexity by approximately 3.8-fold (**Extended Data Fig. 1i-j**). Together, these optimizations ensure robust assay performance across DNA and RNA readouts.

Benchmarking against established standards confirmed the superior performance of ME-seq. In the species-mixing experiment (human HEK293 and mouse NIH/3T3), ME-seq demonstrated accurate species separation, exceptionally low collision rates (<3.1%, **Fig. 1c**), and uniform genomic coverage (**Fig. 1b, c**). Notably, ME-seq outperformed snmCAT-seq, the current leading single-cell methylome-transcriptome assay, in data density, recovering substantially more genes per cell (7,517 vs 4, 522) and comparable or higher numbers of chromatin fragments. Specifically, human cells passing filters contained an average of 87,592 and 50,483 RNA UMIs, 7,517 and 6,272 RNA genes, 37,289 and 1.99M unique DNA fragments, and 73,955 and 1.23M CpGs per cell from Open-ME-seq and Closed-ME-seq, respectively (**Fig. 1b, Extended Data Fig. 1g, Supplementary Table 2**). Compared to the 10x Genomics scATAC-RNA multi-ome assay, Open-ME-seq recovers a comparable number of chromatin fragments, but a significantly higher number of genes (7,517 genes vs 4,631 genes, *p*<2.2×10^−16^, *t*-test). Aggregated ME-seq methylation profiles from as few as 376 cells showed a Pearson correlation >98% with bulk WGBS data, confirming that ME-seq provides bulk-level accuracy at single-cell resolution (**Fig. 1d, e**). Importantly, compared with existing multi-omic platforms, ME-seq reduces library preparation and sequencing costs by orders of magnitude (**Fig. 1f-h, Supplementary Table 3**).

Finally, we demonstrated the versatility of ME-seq by generating the first trimodal single-cell datasets across diverse primary tissues, including human and mouse brains, human fetal liver, and developing mouse embryos (**Fig. 1h, Extended Data Fig. 1k-o, Supplementary Table 4**). Across these tissues, ME-seq consistently captured high-quality, integrated profiles, establishing a robust and democratized framework for dissecting the epigenetic logic of complex biological systems at a scale previously thought unattainable.

### Trimodal atlas charts the cellular and regulatory landscape of aging and Alzheimer’s brain

Having established the robustness of ME-seq, we leveraged its scalability to dissect the temporal and multimodal regulation of AD with a commonly used mouse model of AD amyloid pathology. We profiled the prefrontal cortex (PFC) and hippocampus (HPF), regions highly vulnerable to AD, from C57BL/6 wild-type (WT) and age-matched 5xFAD mice at 3, 6, and 9 months of age (**Fig. 2a**). The 5xFAD model is an early-onset AD model carrying human APP and PSEN1 mutations (APP KM670/671NL (Swedish), APP I716V (Florida), APP V717I (London), PSEN1 M146L, PSEN1 L286V)^24^. This time window captures the critical transition from the initial measurable amyloid burden at 3 months to the onset of memory deficits and profound physiological decline by 9 months.

**Figure 2:**
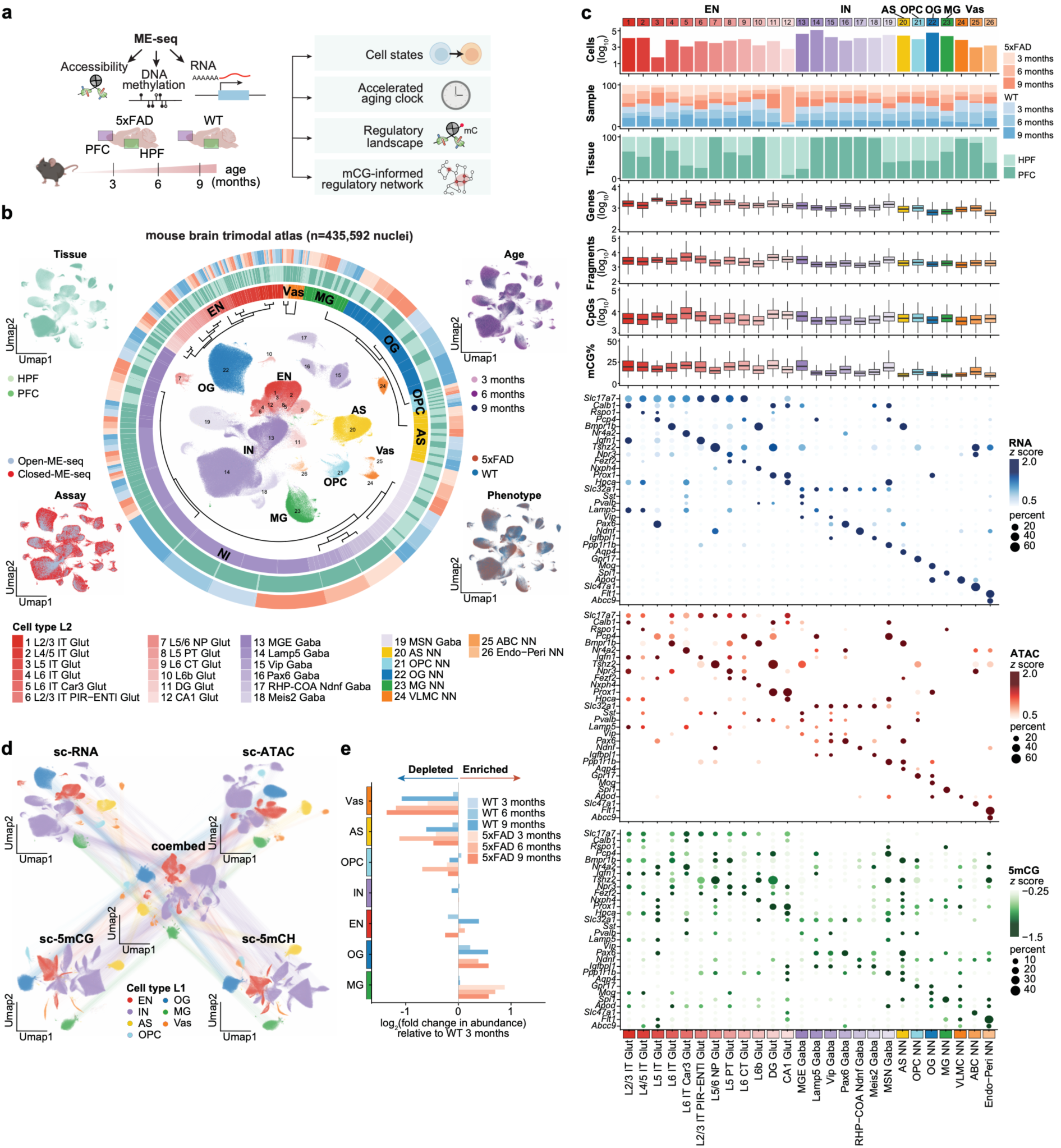
Single-cell trimodal atlas charts the cellular and regulatory landscape of aging and Alzheimer’s brains. (a) Study design using ME-seq to profile wild-type (WT) and 5xFAD mice across the prefrontal cortex (PFC) and hippocampus at 3, 6, and 9 months. (b) UMAP visualizations of the brain atlas derived from the RNA portion of the data with the surrounding hierarchical cellular taxonomy and annotations indicating the cell distribution by cell type, brain region, age, and genotype. Flanking UMAPs show the same embedding colored by tissue region, age, assay type, and genotype. Cell-type annotations and their hierarchical relationships are shown below, with colors corresponding to the clusters in the UMAP. (c) Cluster composition, data quality, and multi-omic characterization of cell populations. Top, bar plots show the number of cells per cluster, the proportion of cells from WT and 5xFAD mice across 3, 6, and 9 months, and the proportion of cells originating from PFC or hippocampus. Middle, box plots summarize data quality metrics across clusters, including the number of detected genes, unique DNA fragments, unique CpG sites, and the average DNA methylation level per cell. Bottom, dot plots display canonical cell-type marker genes, showing RNA gene expression, gene activity scores inferred from chromatin accessibility, and gene body DNA methylation levels. (d) UMAP visualizations of individual data modalities and their joint embedding. All UMAPs are colored by cell-type annotations derived from the scRNA-seq data. Lines connecting the cells indicate correspondence between modality-specific embeddings and the integrated representation. (e) Bar plot showing changes in cell population abundance, normalized to those observed in WT mice at 3 months.

The ME-seq platform demonstrated exceptional technical reproducibility, with near-perfect concordance across replicates (Pearson correlation > 0.99 for all modalities; **Extended Data Fig. 2a**). After stringent filtering, we obtained a dataset of 435,592 high-quality nuclei, representing one of the largest single-cell methylome resources to date (**Fig. 2b; Supplementary Table 5**). Despite the high-throughput nature of the assay, each nucleus yielded high-information content across RNA and DNA (average 2,106 UMIs and 6,950 unique CpGs) at relatively low sequencing depth, underscoring the efficiency of the assay (**Extended Data Fig. 2b**).

Clustering of RNA profiles identified seven major cell types (cell type L1)—excitatory neurons (EN, 15.31%), inhibitory neurons (IN, 55.89%), astrocytes (AS, 5.87%), oligodendrocytes (OG, 13.22%), oligodendrocyte progenitors (OPC, 2.1%), microglia (MG, 5.13%), and vascular cells (Vas, 2.35%)—based on known marker genes (**Fig. 2b-c, Supplementary Table 6**). All major cell types are uniformly detected in Open- and Closed-ME-seq, indicating consistent cell-type representation across assay configurations (**Fig. 2b**). Leveraging the large number of profiled nuclei, ME-seq further resolved fine-grained cellular substructures (cell type L2, **Extended Data Fig. 2c**). For example, excitatory neurons (*Slc17a7*+) were partitioned into 12 glutamatergic subpopulations, such as L2/3 IT Glut (*Calb1*+), L4/5 IT Glut (*Rspo1*+), L5 IT Glut (*Pcp4*+), L6 IT Glut (*Bmpr1b*+), and L6 IT Car3 Glut (*Nr4a2*+) (**Fig. 2c**). Region-specific populations, such as DG and CA1 glutamatergic neurons in the HPF, were detected at expected frequencies, further validating the anatomical fidelity of the atlas^25^.

Across canonical cell type marker genes, gene expression showed a positive correlation with gene activity inferred from chromatin accessibility (Methods) and a negative correlation with gene body DNA methylation (**Fig. 2c, Extended Data Fig. 2d**), consistent with known regulatory relationships. Importantly, independent low-dimensional embeddings of DNA read counts, mCG levels, and mCH (H=A, T, C) levels recapitulated RNA-defined cell types (**Fig. 2d, Extended Data Fig. 2c,d**), demonstrating that each modality independently captures biologically meaningful variation. Notably, independent low-dimensional embeddings of each modality—RNA, ATAC, mCG, and mCH—recapitulated the same cell-type identities, while a Weighted Nearest Neighbor (WNN) integration further sharpened the resolution of cellular diversity (**Fig. 2d**), highlighting the complementary information provided by each modality. Further, cell-type annotations showed strong concordance with a published single-nucleus RNA-seq dataset (**Extended Data Fig. 2e, f**), supporting the accuracy and generalizability of the atlas. Together, these results highlight the scalability of ME-seq to map complex tissues at fine resolution.

The scale of this dataset enabled systematic analysis of how aging and genotype reshape the brain’s cellular composition. While most age-related trends were shared between genotypes (**Fig. 2e**), such as the decline in oligodendrocyte progenitors, astrocytes, and vascular cells and the expansion of mature oligodendrocytes, microglia displayed a distinct, genotype-specific trajectory. Microglia abundance remained stable in WT mice but increased markedly in 5xFAD mice by 3 months, coinciding with the early amyloid deposition, and remained elevated throughout disease progression.

Beyond composition shifts, ME-seq uncovered widespread, modality-specific molecular alterations. We identified thousands of differentially regulated features across all three modalities **(Extended Data Fig. 2h-i, Supplementary Table 7**, FDR<0.05), along with global differences in DNA methylation levels between genotypes (**Extended Data Fig. 2g**). Intriguingly, transcriptomic changes, assessed by differential analysis between age-matched WT and 5xFAD mice, were most pronounced at 3 months, with many differentially expressed genes shared across multiple cell types. In contrast, alterations in chromatin accessibility and DNA methylation were more restricted and highly cell-type specific. For instance, inhibitory neurons showed the most significant epigenetic remodeling at 3 and 6 months, whereas oligodendrocytes exhibited a specific surge in epigenetic shifts at 3 months. This modality-specific timing suggests that the “regulatory pace” of AD is not uniform across cell types, further emphasizing the necessity of trimodal profiling to resolve the latent dynamics of neurodegeneration.

### Accelerated aging in DAM

Given the pronounced and unique expansion of microglia in the 5xFAD model, we performed a high-resolution dissection of microglial heterogeneity and state transitions. Clustering of microglia identified five transcriptionally distinct subsets: homeostatic (*P2ry12*+, *Cx3cr1*+), disease-associated (DAM; *Trem2*+, *Cstb*+), antigen-presenting (MHC; *Cd74*+), cycling/proliferating (*Mcm2*+), and a distinct population of border-associated macrophages (BAM; *Cd163*+) that had been initially annotated as microglia^26^. The microglial landscape was dominated by the transition from homeostatic to DAM states, with the DAM fraction rising from 1.43% at 3 months to over 29.72% by 9 months in the 5xFAD brain, while remaining virtually absent in WT controls (**Fig. 3a, Extended Data Fig. 3a-e**). Notably, integrating multimodal profiles improved sensitivity and interpretability relative to single-modality analysis (**Extended Data Fig. 3b-d**). Differential expression analysis between DAM and homeostatic microglia confirmed robust enrichment of canonical DAM markers, including *Cst7*, *Axl*, *Tmem163*, *Apoe*, and *Lpl*, supporting the accuracy of subtype annotation (**Extended Data Fig. 3f**). Consistent with this progression, a DAM activation score—computed from the expression of 71 established DAM genes^27^—was significantly higher in DAMs at 6 and 9 months compared with homeostatic microglia or DAM at 3 month (**Fig. 3b, Extended Data Fig. 3g**).

**Figure 3:**
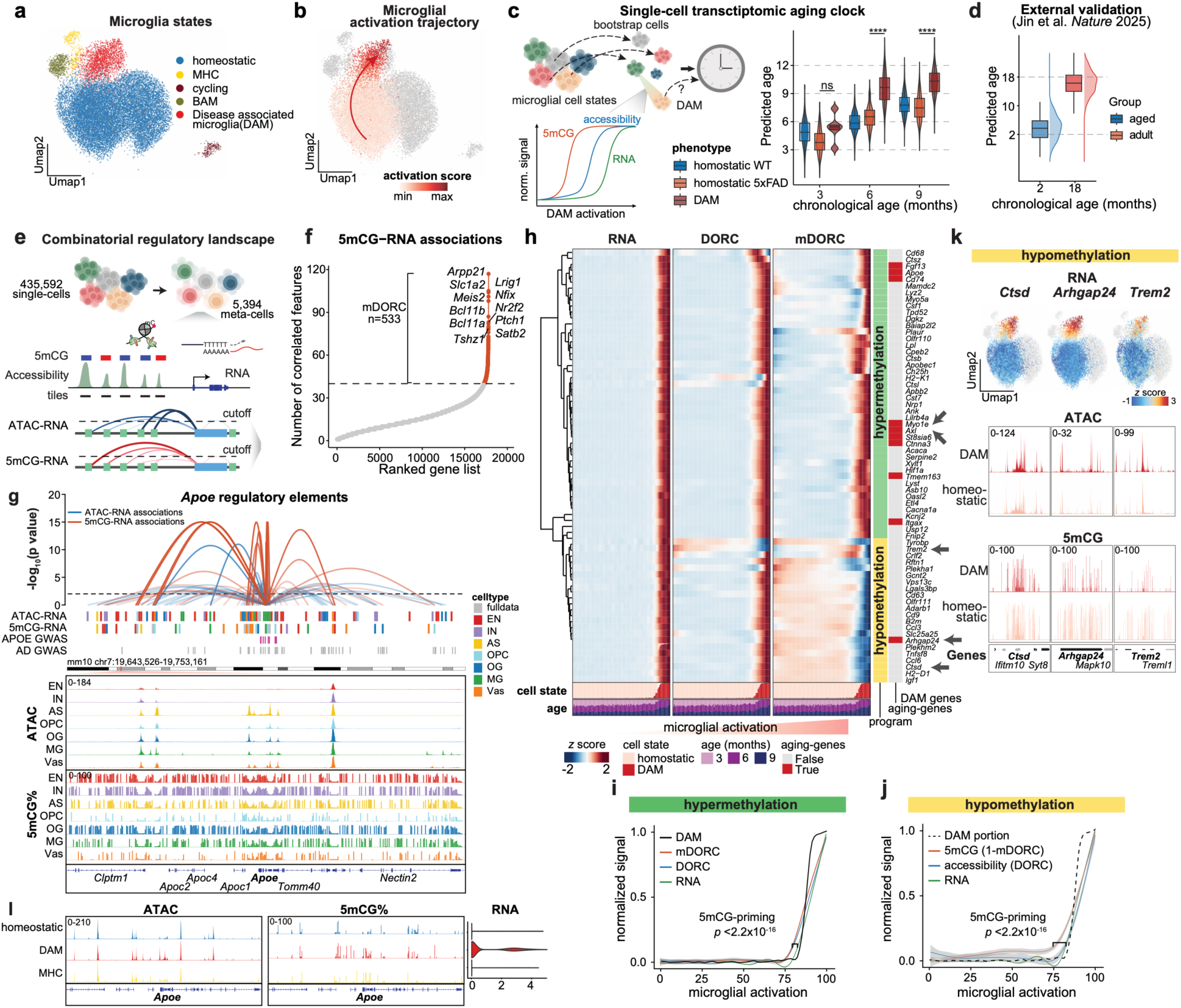
Early microglial priming by DNA methylation. (a) UMAP of microglia colored by subtypes. (b) UMAP of PFC microglia colored by the microglial activation score; arrow indicates increasing activation. (c) Schematic of the aging-clock framework trained on homeostatic microglia and applied to DAM, revealing accelerated biological aging in DAM at 6 and 9 months. (d) Validation of the aging clock using 2- and 18-month-old microglia from Jin et al. *Nature* 2025. Biological ages were predicted using the model trained in this study, demonstrating generalizability. (e) Schematic of the combinatorial CRE analysis framework. Transcriptionally similar cells are aggregated into metacells, and CREs are identified by testing correlations between chromatin accessibility or DNA methylation in genomic tiles and gene expression across metacells. (f) Number of CREs whose DNA methylation levels are significantly correlated (*FDR* < 0.01) with expression of each gene; genes are ranked by the number of associated CREs, with top genes labeled. (g) Genome tracks showing representative CRE-gene associations at the canonical AD marker gene *Apoe*. Loops denote significant CRE-*Apoe* correlations, with loop height indicating significance. Tracks below indicate identified CREs; additional tracks show chromatin accessibility and 5mCG levels by cell type. (h) Heatmaps showing gene expression, DORC scores (mean accessibility of gene-associated CREs), and mDORC scores (mean methylation of gene-associated CREs) across metacells ordered by activation score. Additional below heatmaps indicate the proportion of DAM cells and the corresponding metacell age. (i, j) Temporal ordering of regulatory layers during microglial activation, highlighting 5mCG-mediated priming. Line plots show the mean normalized gene expression, chromatin accessibility, and DNA methylation in hypermethylation (i) and hypomethylation (j) programs. (k) Representative hypomethylation-program genes. UMAPs display normalized RNA expression. Genome tracks show aggregated epigenetic profiles around the corresponding loci. (l) Aggregated accessibility and DNA methylation around the *Apoe* locus, alongside *Apoe* expression.

When examining these transitions, we observed a fundamental divergence between chronological aging and disease progression. In WT mice, microglia aged chronically without ever adopting the DAM phenotype. Conversely, in 5xFAD mice, the emergence of DAMs was not a stochastic response to pathology but followed a progressive trajectory that mirrored, yet outpaced, the normal aging process. This led us to hypothesize that AD pathology acts as a phenotypic accelerator, hijacking intrinsic, age-associated molecular programs to drive microglia into the DAM state. To test this hypothesis and decouple the effects of disease from chronological age, we constructed a microglia-specific aging clock using Lasso regression with bootstrap resampling (**Fig. 3c**, **Extended Data Fig. 3h**). Importantly, the model was trained exclusively on homeostatic microglia—representing the “baseline” of healthy aging. By projecting this baseline model onto the disease-associated states, we could determine if the DAM phenotype represents a *de novo* pathological program or a hyper-accelerated version of the natural aging trajectory.

The results revealed a striking epigenetic age gap in the AD brain. While 6-month-old homeostatic microglia from both WT and 5xFAD mice showed predicted ages consistent with their chronological age (∼6 months), DAM cells exhibited a profound acceleration of biological aging (*p*<2.2×10^−16^, *t-*test). Specifically, in 6-month-old 5xFAD mice, DAM cells were predicted to be 9.54 months old—effectively exhibiting a molecular signature of an aged brain nearly four months ahead of schedule. This trend was sustained at the 9-month timepoint, where DAM cells continued to show the highest biological age (**Fig. 3c**). The model was further validated using an independent dataset^28^, where it accurately distinguished adults from aged microglia and recovering chronological age (**Fig. 3d**).

Examination of “clock-genes” (**Extended Data Fig. 3i, Supplementary Table 8**) revealed DAM selectively amplify genes positively correlated with natural aging, such as *Oas1d* and *Itgax* (*CD11c*) (**Extended Data Fig. 3j**). These clock-associated features were significantly enriched for Gene Oncology (GO) terms related to antigen processing, immune response, and cytokine binding^29^ (**Extended Data Fig. 3k**). This indicates that the DAM state represents a convergence point where disease-specific inflammatory cues accelerate a pre-existing aging-related program, driving microglia toward a “terminally aged” functional state.

Together, these findings establish that the DAM transition is characterized by a coordinated transcriptomic and epigenomic remodeling that drastically accelerates the natural aging process. This suggests that the pathological vulnerability of the aging brain is due to the disease’s ability to “spend” a cell’s epigenetic capital prematurely.

### Early microglial priming by DNA methylation

To dissect the epigenetic hierarchy underlying the microglial transition to the disease-associated state, we sought to identify the CREs that linked to DAM-associated genes and characterize their temporal dynamics. Given that regulatory elements can act over long genomic distances^22^, we developed a combinatorial *cis*-regulatory analysis workflow to systematically and accurately map functional links between DNA methylation and gene expression, as well as between chromatin accessibility and gene expression (**Fig. 3e**). Multiple validation analyses confirmed the robustness of this approach (**Extended Data Fig. 3n-p**). This framework enables systematic, cell-type–resolved identification of functional regulatory elements and their temporal hierarchy, uncovering regulatory relationships that are inaccessible to single-modality or bulk approaches.

By aggregating transcriptionally similar cells into 5,394 metacells^30^ (average 80.7 cells per metacell; **Extended Data Fig. 3l, m**) to overcome sparsity, we identified a high-confidence regulatory landscape consisting of 498,809 ATAC-RNA associations and 286,145 5mCG-RNA associations, in a total of 621,053 CRE-gene pairs (within ± 50kb around TSSs, FDR < 0.01; **Fig. 3f, Supplementary Table 9,10**). Notably, these two regulatory layers showed remarkably limited overlap, suggesting that DNA methylation and chromatin accessibility govern distinct genomic territories. Our framework successfully captured the regulatory logic of AD-associated GWAS variants, including those at the APOE locus. By incorporating the methylome, we linked several distal risk variants to APOE that were not identified in previous GWAS studies, demonstrating that DNA methylation provides an essential, previously “invisible” layer of disease-relevant regulation (**Fig. 3g**).

The scale of this map revealed Domains of Regulatory Chromatin (DORCs)^22^, a subset of genes that were associated with an unusually large number of CREs (*p*<2.2×10^−16^, permutation test) for both DNA methylation and chromatin accessibility. We defined 870 accessibility-driven DORCs (>80 ATAC-RNA associations, **Extended Data Fig. 3o**) and 533 methylation-associated DORC genes (mDORCs; >40 5mCG-RNA associations; **Fig. 3f**). While these sets converge (*p* < 10^−409^; hypergeometric test) on key microglial lineage regulators such as *Spi1* (PU.1) and *Smad4*, the underlying individual CREs were largely modality-specific, suggesting a coordinated but independent dual-layer control of cell fate (**Extended Data Fig. 3q**).

By mapping these associations along the DAM activation trajectory, we uncovered a phenomenon of epigenetic priming: DNA methylation remodeling systematically precedes changes in both chromatin accessibility and transcription (*p*<10^−16^, KS test). We identified two distinct priming programs: a hypermethylation program and a hypomethylation program (**Fig. 3h**). In the hypermethylation program, genes involved in lipid metabolism and metabolic sensing, such as *Apoe*, *Myo1e*, and *Axl* (**Extended Data Fig. 3r)**, undergo early increases in both 5mCG and accessibility that prime the locus for massive transcriptional upregulation (**Fig. 3i**). Conversely, genes in the hypomethylation program, *Ctsd*^31^, *Arhgap24*^32^, and *Trem2*^33^, exhibited early DNA demethylation, followed by accessibility changes, and subsequent transcriptomic onset (**Fig. 3j, k**). These two programs likely reflect the complex genomic context of methylation, where promoter hypomethylation and gene-body hypermethylation both serve to establish “transcriptional competence.” In both instances, the methylome acts as a “latent blueprint,” pre-configuring the genome’s response to amyloid pathology before the first DAM-associated transcripts are synthesized.

Visualization of these methylation-sensitive CREs revealed that they are enriched for CpG-dense regions and uniquely harbor motifs for IRF1 and PU.1 (**Extended Data Fig. 3s-v**). Collectively, these findings establish DNA methylation as an early, modality-specific regulatory layer that sets the trajectory for subsequent chromatin reorganization. By the time accessibility changes are detectable by standard assays, the cell’s pathological fate has already been “locked in” by the methylome.

### Chromatin state reconfiguration during microglial priming

To move beyond locus-specific observations and investigate the systemic mechanisms of DAM priming, we characterized genome-wide chromatin-state dynamics during aging and AD progression. We reasoned that the joint measurement of RNA expression, chromatin accessibility, and DNA methylation could reveal combinatorial regulatory state shifts underlying microglia activation and reveal regulatory programs that are “invisible” to single-modality analyses. To resolve these dynamics, we developed meta-ChromHMM, a framework that integrates all three modalities to infer genome-wide chromatin states at metacell resolution using a hidden Markov model^34^ (**Fig. 4a**). Notably, metacell aggregation reduced sparsity to levels comparable to cell-type-specific pseudobulk profiles (**Extended Data Fig. 4a**). Unlike traditional pseudobulk approaches, meta-ChromHMM preserves the continuity of single-cell trajectories, enabling the detection of disease-specific chromatin reconfigurations that would otherwise be masked by cellular averaging.

**Figure 4:**
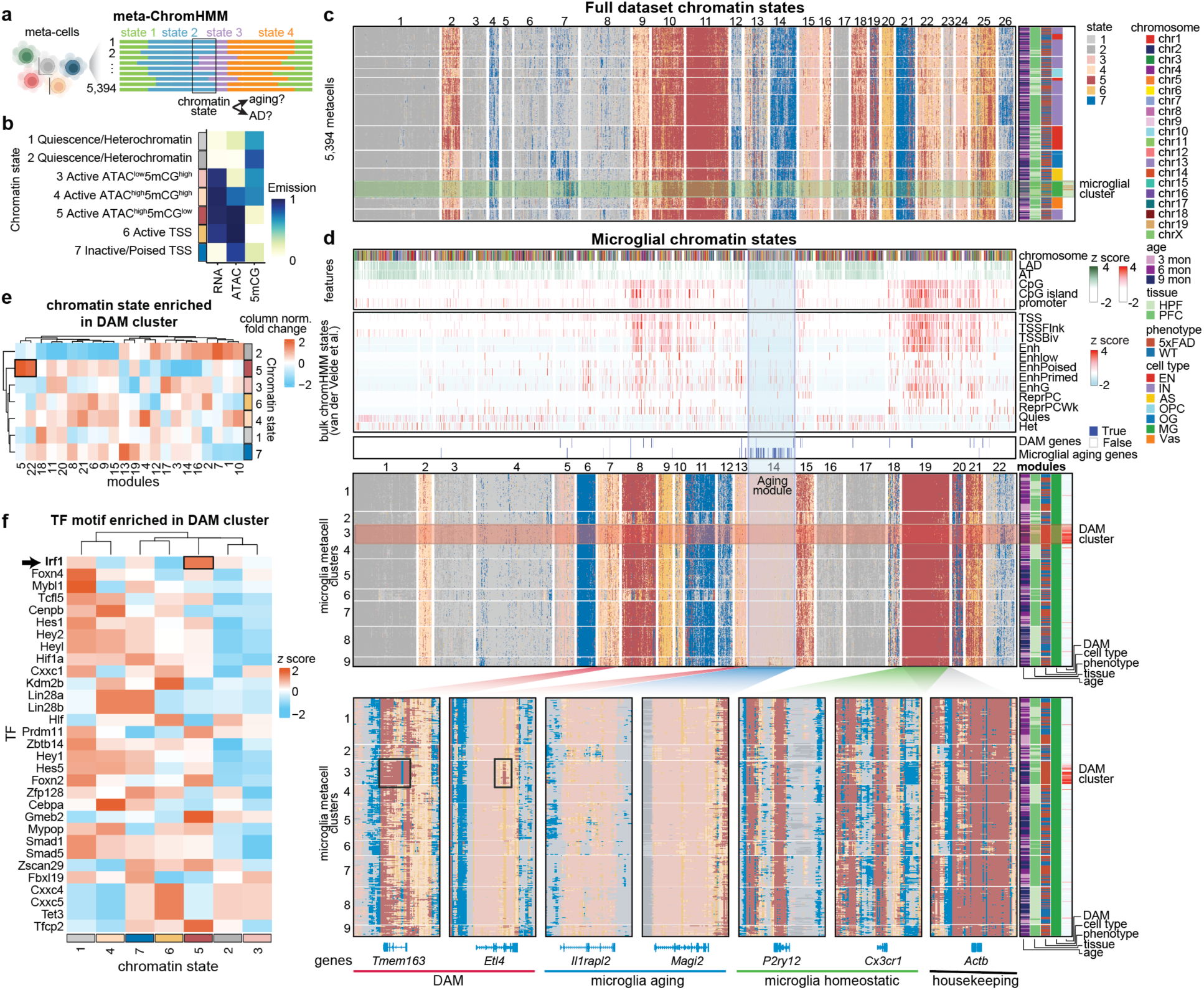
Chromatin state reconfiguration during DAM activation. (a) Schematic illustrating the meta-ChromHMM framework. Metacells are used to generate binarized input matrices from gene expression, chromatin accessibility, and DNA methylation signals across fixed-length genomic windows, which are then integrated to infer genome-wide chromatin states. (b) Heatmap showing emission probabilities for each modality across the seven chromatin states inferred by meta-ChromHMM. Annotations indicate the functional interpretation of each chromatin state. (c) Heatmap showing the distribution of chromatin states inferred from the full dataset across seven major cell types, two brain regions, three ages, and genotypes. 111,972 genome-wide 25-kb bins are grouped into 26 clusters. Metacell clusters enriched for microglia are highlighted. (d) Heatmap showing the distribution of chromatin states inferred from microglia. Clusters enriched for DAM and clusters enriched for microglial aging gene programs are highlighted. Annotations above the heatmap indicate CpG density, A/T content, lamina-associated domain (LAD) enrichment, and publicly reported EpiMap chromatin states. (e) Heatmap showing enrichment of each chromatin state in DAM-enriched clusters compared with other metacell clusters, calculated using the genome-wide bin clustering shown in panel (d). (f) Heatmap showing enrichment of the top 30 most variable TF motifs across the seven chromatin states within DAM-enriched clusters.

Systematic optimization (**Extended Data Fig. 4a-f**) revealed that a seven-state model maximized resolution and interpretability across 435,592 nuclei, identifying distinct functional compartments including active TSS, poised/inactive TSS, and three transcriptionally active gene-body states (active ATAC^high^5mCG^low^, active ATAC^high^5mCG^high^, active ATAC^low^5mCG^high^; **Fig. 4b**). These active states were distinguished primarily by their DNA methylation levels rather than accessibility alone. Validation against global nuclear architecture, including CpG density, A/T content, lamina-associated domains (LADs), and EpiMap annotations^35^, confirmed that meta-ChromHMM accurately captures the higher-order organization of the genome^36^ (**Fig. 4c; Extended Data Fig. 4g**).

Focusing on microglia, we identified a profound reorganization of the chromatin landscape as cells transition toward the DAM state (**Fig. 4d**). We identified distinct chromatin modules for natural aging (Module 14, odds ratio = 37.35, *p* < 0.001) versus AD progression (Modules 5 and 22; **Fig. 4e**). Module 14 was characterized by high A/T content and enrichment of LAD. Intriguingly, canonical DAM genes, such as *Tmem163* and *Etl4*, transitioned from an active, highly methylated state to an active, low-methylation state without substantial changes in chromatin accessibility. This finding suggests that for a significant subset of the AD-associated genome, DNA methylation remodeling—rather than chromatin opening—is the dominant architect of cellular activation. This large-scale reorganization across broad genomic regions implies that microglial activation is not driven by isolated regulatory events, but by a coordinated “epigenetic switch” that repositions the genome for rapid inflammatory response.

Crucially, we found that the DAM-associated state (ATAC^high^/5mCG^low^) was selectively enriched for the motifs such as IRF1 relative to all other microglial clusters (**Fig. 4f**). IRF1 is a master regulator of interferon-driven transcriptional networks and is known to coordinate the magnitude of microglial reactive states^37^. While IRF1 has been conventionally studied through changes in expression or accessibility, our data suggest a more nuanced “gating” mechanism. We observe that IRF1 motifs are frequently embedded within regulatory elements that undergo demethylation while remaining accessible^38^, suggesting that DNA methylation acts as a biochemical gate that controls IRF1 engagement.

Collectively, these genome-wide analyses reveal that DAM priming is governed by a large-scale reconfiguration of the chromatin landscape, initiated by DNA methylation shifts that systematically precede accessibility changes. These patterns, which identify the methylome as the primary initiator of the microglial “epigenetic terminal state,” were only discernible through the trimodal integration enabled by ME-seq. These results establish DNA methylation as the central regulatory axis of microglial activation, providing a new paradigm for how the genome records and responds to neurodegenerative stimuli.

### IRF1 is a methylation-sensitive master regulator that primes microglial activation

We next sought to identify the TFs that mediate DNA methylation-dependent DAM activation. ME-seq data provides a unique opportunity to infer the TF binding activity influenced by local DNA methylation states at single-cell resolution. We developed a computational framework to quantify DNA methylation–associated TF motif activity data (Methods), where high scores reflect increased methylation levels at a given TF’s binding sites (**Fig. 5a; Extended Data Fig. 5a, b**). This metric allows us to capture the regulatory potential—or “licensing“—of a TF independently of its current occupancy or expression level.

**Figure 5:**
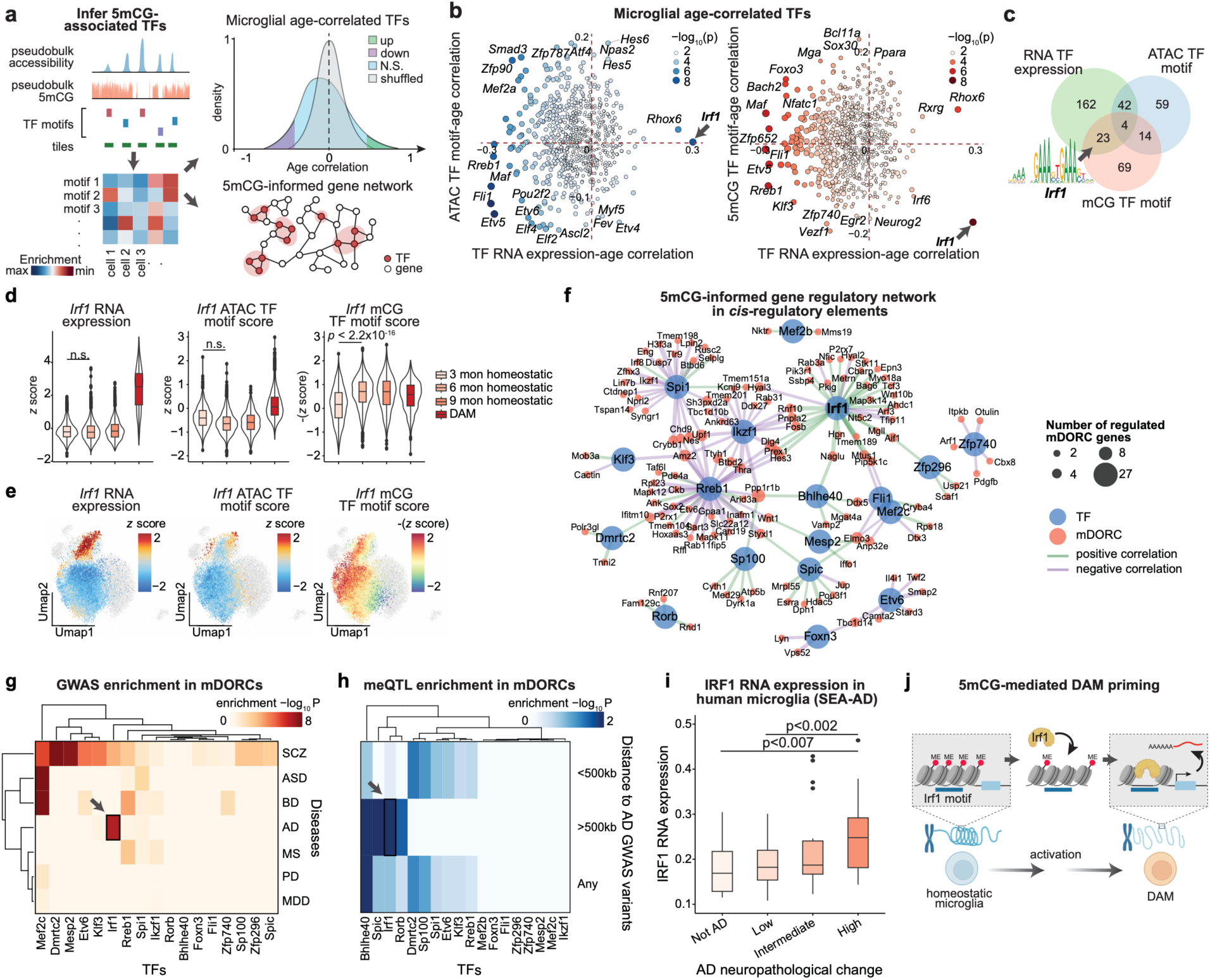
IRF1 is a key TF in microglia aging and primes DAM activation. (a) Workflow for inferring TF motif activity from chromatin accessibility or DNA methylation across 500-bp windows in single cells or metacells. Motif activity scores are computed relative to a matched null distribution, correlated with age, and integrated with *cis*-regulatory analyses to identify TF drivers and regulatory networks. (b) Age-correlated TF activities inferred from different regulatory modalities in microglial metacells. Each dot represents a TF; x-axis, correlation between TF RNA expression and age; y-axis, correlation between ATAC-based (left) or 5mCG-based (right) motif activity and age. (c) Overlap of TFs whose activities are significantly age-associated across modalities, highlighting *Irf1* RNA expression and 5mCG-inferred motif activity, but not ATAC-inferred activity. (d) Violin plots of *Irf1* expression (left), ATAC-inferred motif activity (middle), and 5mCG-inferred motif activity (right) in homeostatic microglia at 3, 6, and 9 months and in DAM. (e) UMAPs of microglia colored by *Irf1* expression (left), ATAC-inferred motif activity (middle), and 5mCG-inferred motif activity (right). (f) 5mCG-informed TF regulatory network linking TF expression to mDORC scores; edge width reflects significance (−log_10_ FDR). *Irf1* is highlighted as a central regulator. (g) Heatmap depicting the enrichment of public disease-associated GWAS variants within TF target gene–associated mDORCs. Brain disorders include AD, PD, multiple sclerosis (MS), major depressive disorder (MDD), bipolar disorder (BD), and schizophrenia (SCZ). (h) Heatmap depicting the enrichment of AD-associated CpG sites within TF target gene–associated mDORCs. (i) IRF1 gene expression levels in human microglia across different groups from a previously published study (SEA-AD). (j) Proposed model linking DNA methylation to *Irf1* activity and microglial activation. Schematic illustrating a putative mechanism by which hypomethylation at *Irf1* binding sites enhances *Irf1* TF binding activity, leading to subsequent chromatin remodeling and transcription activation, and primes the transition from homeostatic microglia to DAM during AD progression.

By comparing motif activity derived from both chromatin accessibility (ATAC) and DNA methylation (5mCG), we identified IRF1 as a uniquely microglia-specific regulator. While ATAC-based IRF1 motif activity was broadly detected across several brain cell types, methylation remodeling at IRF1 sites was exclusive to the microglial lineage (**Extended Data Fig. 5c, d**). Notably, IRF1 RNA expression showed the strongest positive correlation with age among all TFs (*p* = 3.0×10^−9^), yet its chromatin motif activity remained largely unchanged across ages (Pearson r = 0.0008, *p* = 0.49). In contrast, the IRF1 5mCG motif activity score showed a profound negative correlation with age (**Fig. 5b**). Given that IRF1 preferentially binds hypomethylated CpG-containing motifs^39^ (**Fig. 5c**), these results suggest that aging-associated demethylation enhances IRF1 binding capacity independently of chromatin opening.

The temporal resolution of ME-seq revealed a striking “priming” signature during early DAM activation. In the 5xFAD brain, the IRF1 5mCG motif activity was significantly elevated as early as 3 months (**Fig. 5d, e**), preceding detectable changes in IRF1 RNA expression or ATAC motif activity. This suggests that IRF1 functions as an early priming factor, whose regulatory potential is unmasked by DNA methylation loss well before the cell commits to a reactive transcriptional state.

To define the global impact of this mechanism, we constructed a TF-regulatory network by integrating 5mCG TF binding activities, CRE-gene linkages, and transcriptional outputs (Methods). IRF1 emerged as a central hub of microglial regulome, coordinating programs for activation, migration, and inflammatory response^40,41^ (**Fig. 5f**). This network also incorporated established AD regulators^42,43^, including MEF2C, BHLHE40/41, and SPI1 (PU.1), but positioned IRF1 as the primary driver of methylation-mediated transitions (**Fig. 5f; Extended Data Fig. 5e, f**).

To establish the clinical relevance of this axis, we intersected our IRF1-target networks with human genetic data (**Fig. 5g,h**; **Extended Data Fig. 5g,h**). AD-associated GWAS variants (*p* <10^−7^, *z* test; **Fig. 5g**) and methylation quantitative trait loci (meQTLs; **Fig. 5h**) were significantly enriched within IRF1-associated 5mCG-RNA loops, but not ATAC-RNA loops (**Extended Data Fig. 5h**). Critically, we found that IRF1 target loops were enriched for distal CpG sites linked to AD risk that are otherwise “invisible” to standard GWAS mapping (**Fig. 5h**). Finally, we confirmed that IRF1 upregulation is not limited to mouse models; IRF1 is significantly overexpressed in human AD microglia^44^, suggesting a conserved role in human neurodegeneration (**Fig. 5i**).

Together, these results identify IRF1 as the central “gatekeeper” of microglial aging. By demonstrating that demethylation at IRF1 binding sites constitutes the earliest measurable event in the DAM transition, we establish DNA methylation remodeling as the critical “license” that enables inflammatory transcriptional programs to manifest during AD progression (**Fig. 5j**).

## Discussion

Single-cell profiling technologies have transformed our ability to define cellular identities, yet our view of the regulatory hierarchies in the aging brain has remained incomplete. By omitting the DNA methylome—the genome’s most stable and heritable epigenetic repository—previous multi-omic studies have largely focused on the “active” execution of gene programs, remaining blind to the latent regulatory memory that precedes pathological activation. Here, we overcome the technical and economic barriers of single-cell methylomics through ME-seq. By achieving a 100-fold reduction in cost and enabling trimodal integration, ME-seq transitions the field from descriptive atlases to a predictive framework for understanding the epigenetic temporal hierarchy of neurodegeneration.

Our time-course analysis of the 5xFAD model reveals a striking alignment between epigenetic remodeling and the progression of Alzheimer’s pathology. In the 5xFAD brain, amyloid-beta (Aβ) deposition begins as early as 1.5-2 months, reaching a critical threshold by 3 months. We observe that this initial amyloid insult coincides with the earliest detectable DNA methylation shifts in homeostatic microglia. Crucially, these alterations do not merely mirror existing pathology but act as a foundational priming layer that precedes overt DAM transition. This finding extends the current understanding of the “amyloid cascade” by suggesting that Aβ acts as a catalyst for epigenetic reprogramming, establishing a state of transcriptional competence long before the peak of neuroinflammation.

A central discovery of this study is the identification of IRF1 as a master methylation-sensitive “gatekeeper” of microglial activation, along with several candidate master regulators of DAM priming^42,43^. While previous studies have implicated IRF1 in regulating the expression of core DAM genes like *Apoe*, *Axl*, and *Trem2*^37^, and established its requirement for IRF8-mediated *Il1b* expression in reactive microglia^45^, our data provide the missing regulatory link. We propose a model where early Aβ deposition triggers DNA demethylation at IRF1-sensitive enhancers, lowering the kinetic barrier for interferon-responsive programs. By 6 and 9 months, as plaque burden intensifies and the DAM population expands, this primed IRF1 axis drives a self-amplifying inflammatory cycle. In this cascade, Aβ-induced IL-1β further stimulates *Irf1* expression through NF-κB and MAPK signaling, cementing the pathological state and accelerating the transition from homeostatic to reactive phenotypes.

This regulatory cascade effectively blurs the traditional distinction between “normal aging” and “disease progression.” Our single-cell aging-clock analysis reveals that DAM cells exhibit a marked acceleration of biological age compared to their homeostatic counterparts. This suggests that AD pathology does not simply induce a *de novo* disease program but rather hijacks intrinsic age-associated molecular programs—specifically those governed by DNA methylation drift—to drive cellular exhaustion. The DAM state may thus represent an “epigenetic terminal state,” where the loss of homeostatic methylation signatures renders microglia hyper-responsive to neurodegenerative cues.

Our combinatorial *cis*-regulatory analysis further identifies a subset of methylation-associated regulatory elements that are functionally “invisible” to standard ATAC-seq assays. By 6 months, these CREs show profound methylation changes despite remaining in an inaccessible chromatin state, indicating that the methylome acts as a “latent blueprint” for the neurodegenerative transitions that manifest more fully at 9 months. These findings shift the prevailing view of microglial activation from a chromatin-opening–driven process to one initiated by epigenetic hysteresis, where the methylome records the amyloid insult and “locks” the cell into a pro-inflammatory trajectory.

Despite these advances, several limitations remain. While the 5xFAD model provides a robust scaffold for amyloid-driven changes, the “epigenetic drift” in late-onset AD (LOAD) likely involves broader environmental and genetic interactions. Future applications of ME-seq to human longitudinal cohorts will be essential to validate the IRF1-priming axis in clinical contexts and to investigate how the dosage of AD-associated SNPs correlates with methylation status. Furthermore, while we establish strong associations between methylation remodeling and DAM activation, direct causal validation via targeted epigenetic editing—“resetting” the microglial clock—will be the next frontier in therapeutic development.

In conclusion, ME-seq offers a democratized platform to decode the “latent” regulatory states of the genome. By defining the temporal hierarchy where amyloid deposition leads to DNA methylation remodeling and subsequent IRF1-driven microglial activation, we identify a critical window for intervention before irreversible transcriptional and cognitive decline occur. These findings redefine the temporal hierarchy of regulatory events in neurodegeneration, establishing DNA methylation as the primary initiator of pathological state transitions.

Looking forward, ME-seq provides a broadly accessible platform for dissecting epigenetic regulation across development, aging, and disease at the tissue scale. Coupled with perturbative approaches and longitudinal sampling, this technology offers a powerful framework to unravel how DNA methylation encodes cellular memory, constrains plasticity, and drives pathological state transitions. More broadly, the trimodal atlas generated here serves as a foundational resource for understanding the epigenetic logic of brain aging and AD.

## Extended Data

**Extended Data Figure 1:**
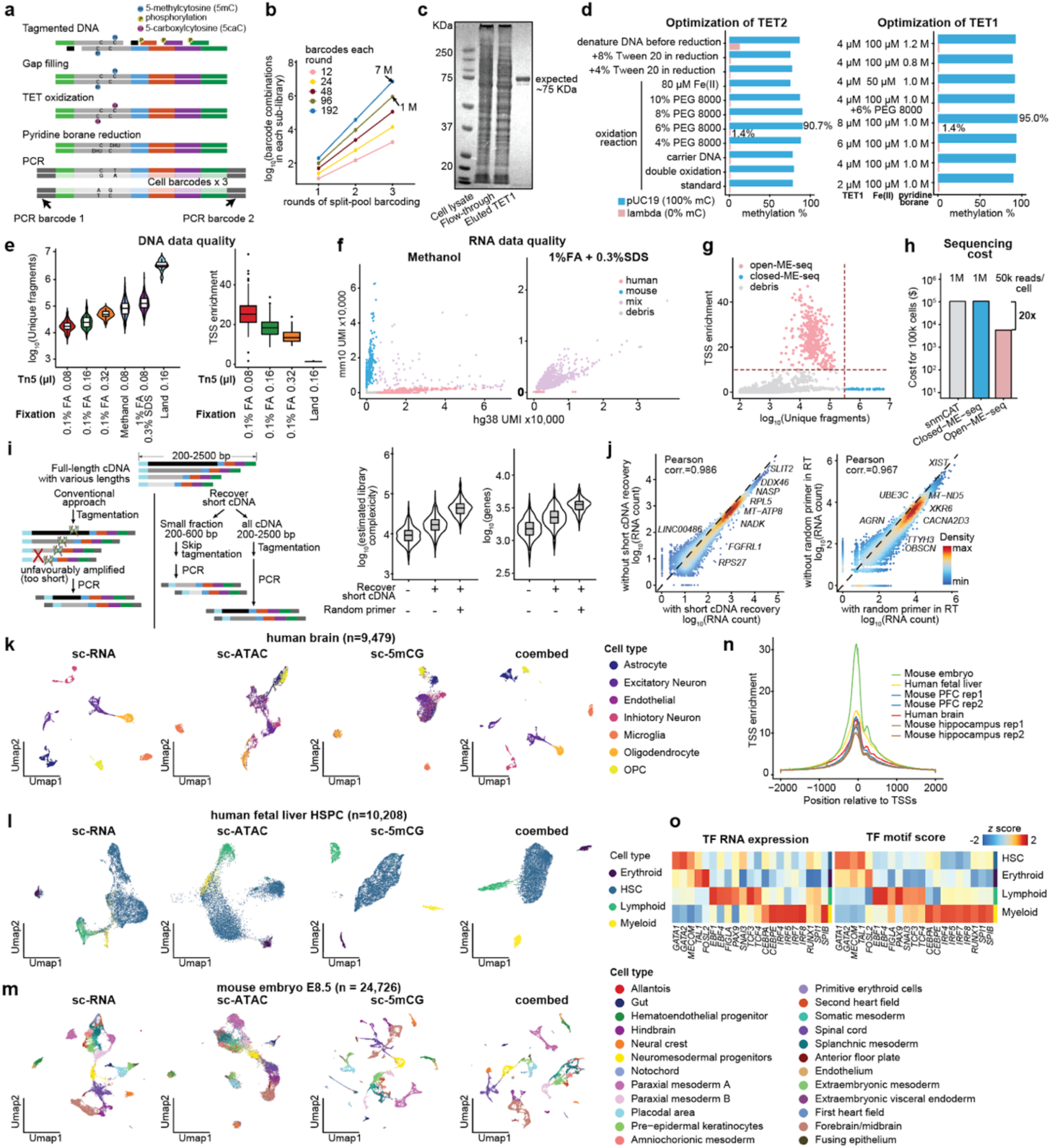
The principle of ME-seq, assay optimization, and data quality control on tissue datasets (related to Fig. 1). (a) The structure of the DNA sequencing library. (b) The expected number of barcode combinations exponentially scales with the rounds of barcoding. (c) SDS-PAGE gel showing the size of the purified TET1 protein. (d) Optimization of DNA methylation profiling using TET1-and TET2-based routes, including oxidation and reduction steps. (e) Optimization of DNA library preparation based on unique DNA fragments per cell, including nucleosome removal methods and Tn5 transposase dosage. (f) Results of cross-contamination tests using cells treated with methanol or with a combination of 1% formaldehyde (FA) and 0.3% SDS. (g) Scatter plot showing the relationship between the number of unique genomic fragments per cell (x-axis) and TSS enrichment (y-axis). Each dot represents a single cell. (h) Bar plot showing the sequencing cost per 100,000 single-cell libraries for snmCAT, Closed-ME-seq, and Open-ME-seq. (i) Schematic showing strategies to improve RNA library quality by short cDNA recovery and random priming during RT, with violin plots illustrating the resulting improvements. (j) Scatter plots comparing gene-level read counts between libraries with and without short cDNA recovery (left) or random priming during RT (right), with libraries downsampled to matched cell numbers and sequencing depth. (k-m) UMAP visualizations of ME-seq across multiple primary tissues, shown from left to right as scRNA-seq, scATAC-seq, sc5mCG, and the co-embedded integration of all modalities. Panels show human brain samples (**k**), human fetal liver HSPC (**l**), and mouse embryo samples (**m**). (n) Enrichment of aggregated reads across ±2 kb of TSSs for mouse embryo, human fetal liver, mouse brain, and human brain libraries. (o) Heatmaps showing gene expression of cell-type—specific TFs (left) and the corresponding TF motif activity scores (right) from the human fetal liver study.

**Extended Data Figure 2:**
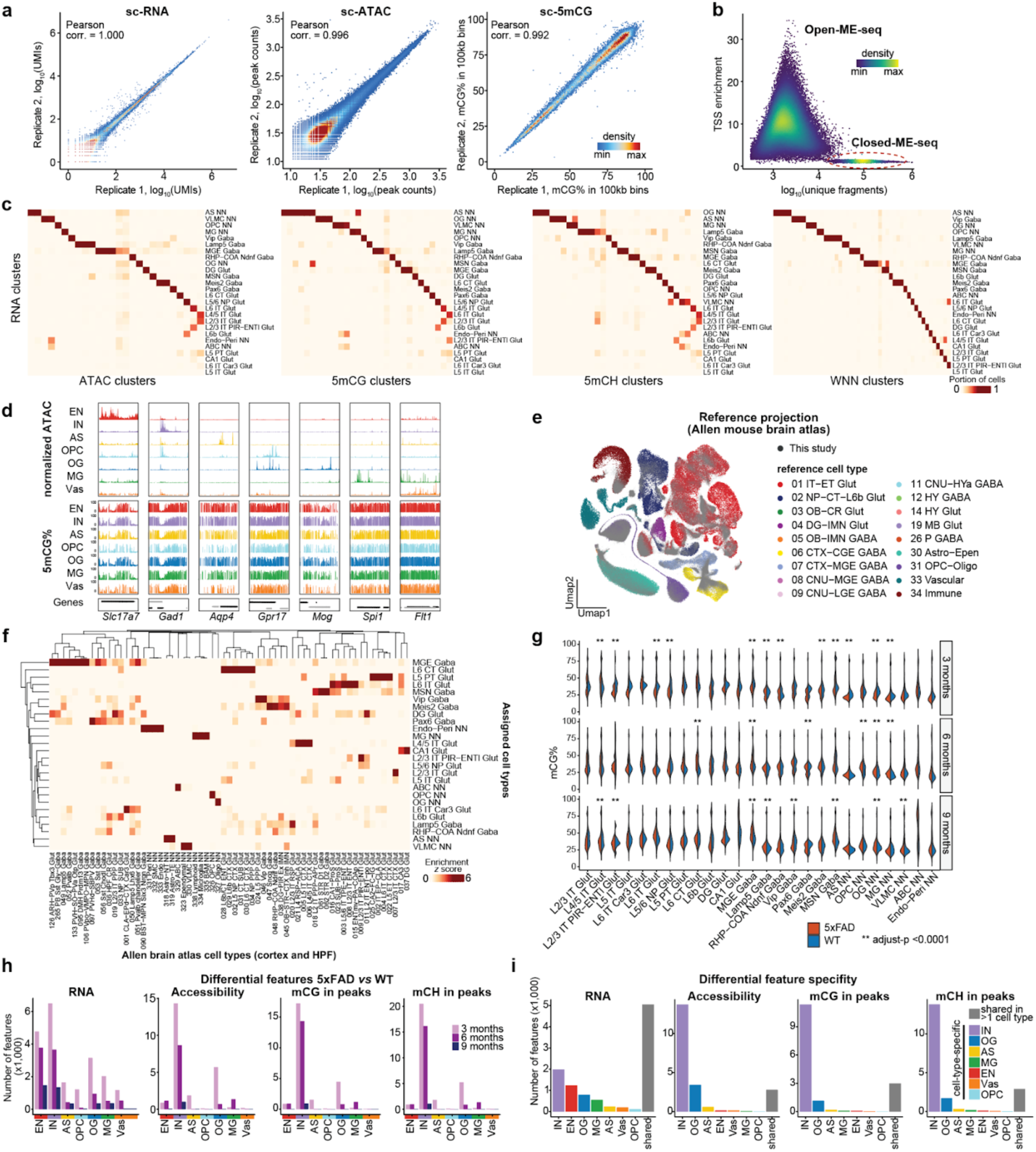
High-quality ME-seq mouse brain data reveals broad congruence across modalities alongside widespread modality-specific differences (related to Fig. 2). (a) Reproducibility of ME-seq profiles between technical replicates. (b) Scatter plot showing transcription start site (TSS) enrichment versus the number of unique fragments per cell for mouse brain data. (c) Heatmap showing the proportion of cells in each RNA-defined cluster that overlap with ATAC-seq, 5mCG, 5mCH, or joint clusters in the mouse brain. (d) Aggregated single-cell tracks of chromatin accessibility and 5mCG at marker genes for each major cell type. (e) UMAP visualization showing reference projection of cells from this study onto the Allen mouse brain atlas dataset (PMC11798837). Colors indicate cell-type annotations from the reference dataset; grey denotes cells from this study. (f) Heatmap showing enrichment of cells from this study overlapping with cell types in the reference dataset. (g) Violin plots showing cell-type-specific differences in global 5mCG levels. ** indicates FDR < 0.0001. (h) Number of differential features identified between 5xFAD and WT mice for each cell type and age (Wilcoxon rank-sum test, FDR < 0.05). (i) Bar plot showing the number of features in (h) that are cell-type-specific or shared across cell types (grey).

**Extended Data Figure 3:**
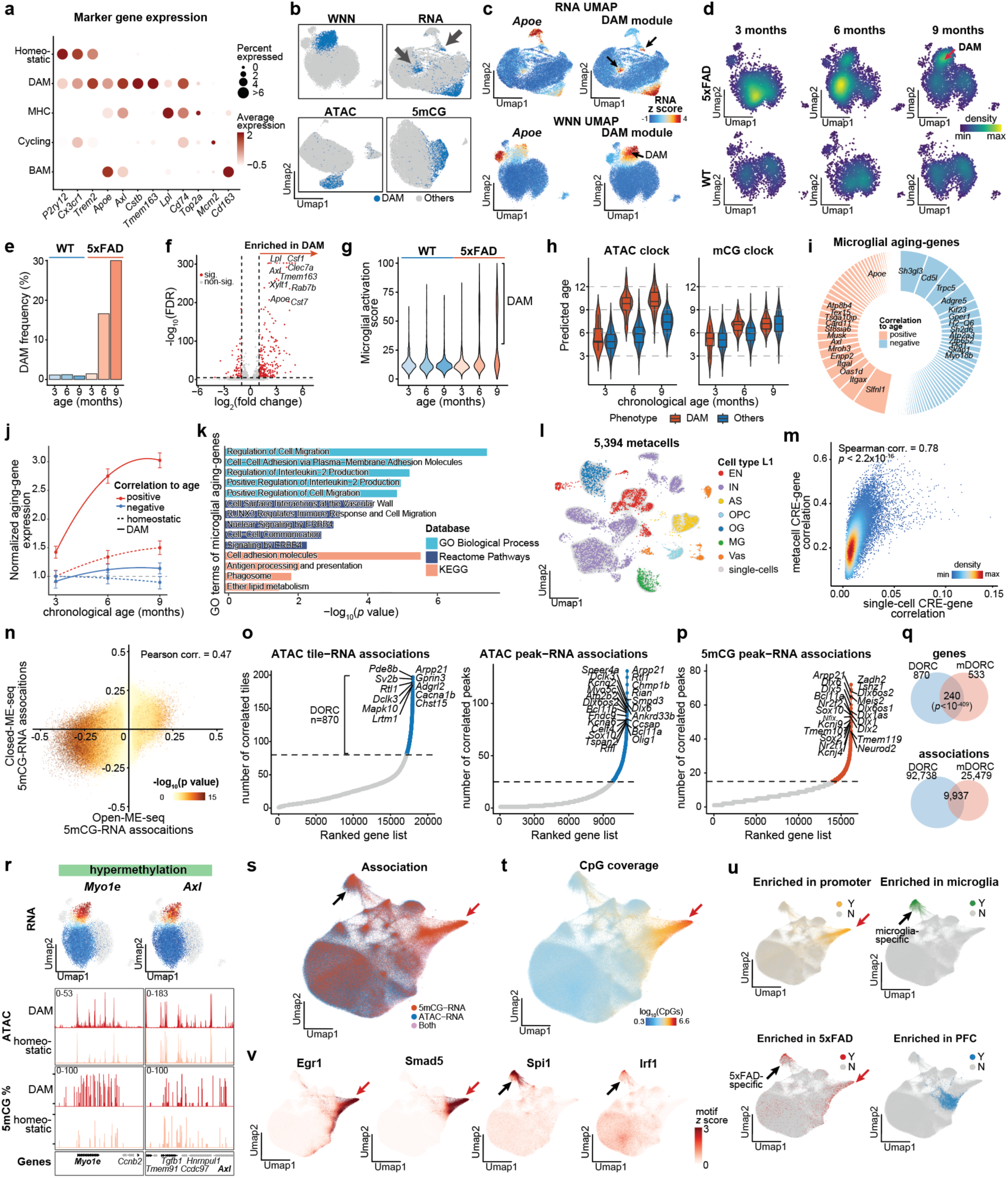
Combinatorial *cis*-regulation analysis identifies disease-specific regulatory elements in microglia subsets (related to Fig. 3). (a) Dot plot showing expression of marker genes across microglial states. (b) Comparison of UMAP visualizations derived from individual data modalities or joint integration. Blue indicates annotated DAM cells. (c) UMAP visualizations showing *Apoe* RNA expression and DAM module scores, calculated from 74 previously reported DAM marker genes. (d) Shifts in microglial cell states during aging, highlighting expansion of DAM cells at 6 and 9 months. (e) Quantification of changes in the frequency of annotated DAM cells across ages. (f) Volcano plot showing differentially expressed genes between DAM and homeostatic microglia. Genes most strongly enriched in DAM are labeled and overlap with known DAM markers. (g) Violin plot showing the distribution of DAM scores across microglial cells. (h) Predicted biological age of WT and 5xFAD microglia using aging clocks derived from chromatin accessibility and DNA methylation. (i) Pie chart showing the relative contribution (weights) of genes to the transcriptomic aging clock. (j) Average RNA expression changes of genes contributing to the transcriptomic aging clock, highlighting significant upregulation of genes positively correlated with age in DAM. (k) Gene Ontology (GO) term enrichment analysis of genes contributing to the transcriptomic aging clock. (l) UMAP of the mouse brain dataset highlighting uniformly distributed metacells defined from scRNA-seq. (m) Spearman correlations between metacell-aggregated or single-cell gene expression and DNA methylation. (n) Comparison of Spearman correlations between RNA expression and DNA methylation at CREs. (o) Number of genomic tiles (left) or open chromatin peaks (right) whose accessibility is significantly correlated (FDR < 0.01) with expression of each gene. (p) Number of peaks whose DNA methylation levels are significantly correlated (FDR < 0.01) with expression of each gene. (q) Venn diagrams showing overlap between DORC and mDORC genes (top) and between DORC- and mDORC-associated CREs (bottom). (r) Representative examples of hypermethylation-program genes (*Myo1e* and *Axl*). Top: UMAPs showing single cells colored by normalized RNA expression. Middle and bottom: genome tracks showing aggregated chromatin accessibility and DNA methylation, respectively, for DAM and homeostatic microglia at the corresponding loci. (s-u) UMAP visualizations of 621,053 CRE-gene pairs inferred from the full dataset, colored by associated modality (s), CpG coverage within the CRE (t), and enrichment across genomic regions, cell types, genotypes, brain regions (u), and TF motif representation (v).

**Extended Data Figure 4:**
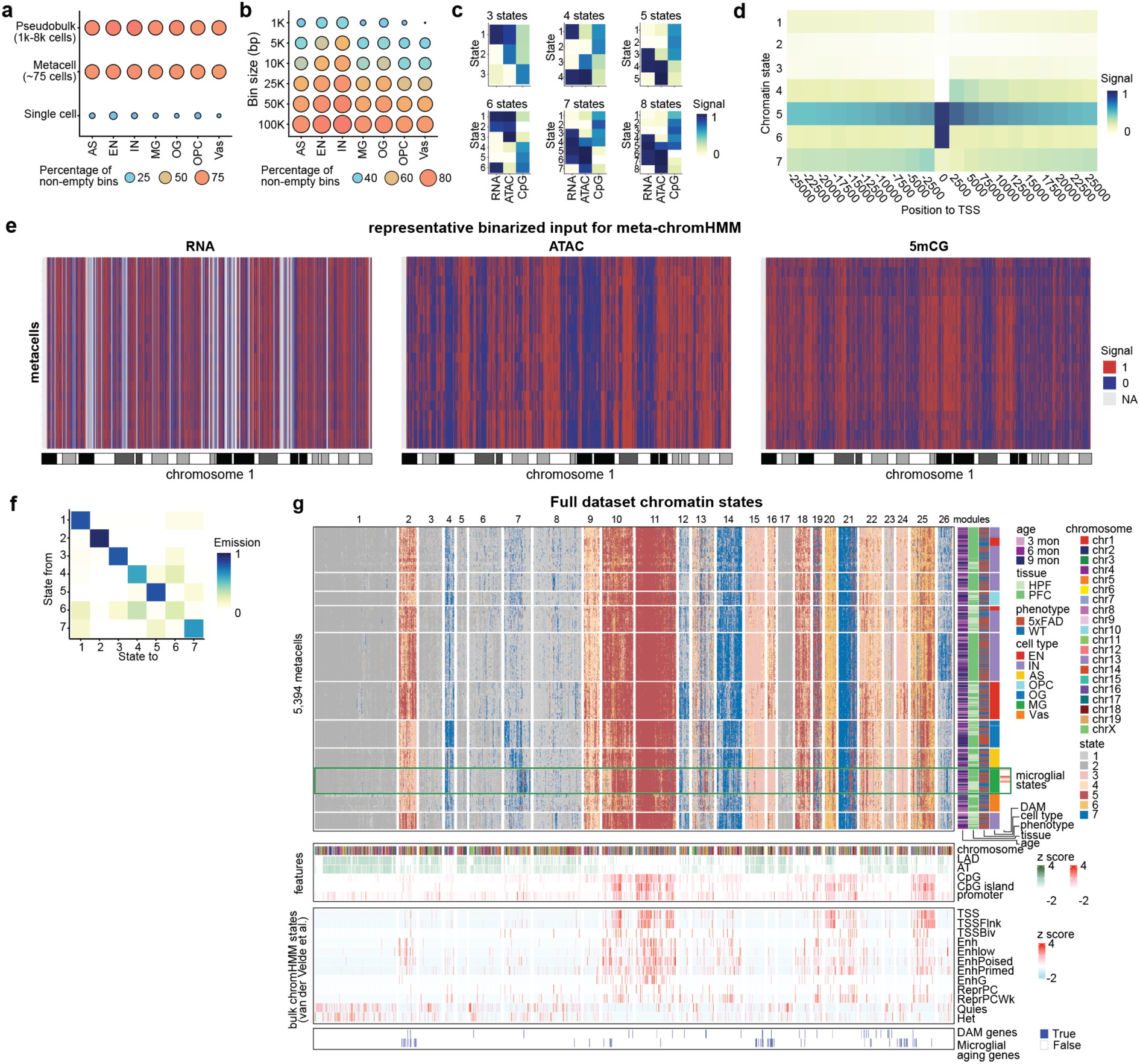
Optimization of meta-ChromHMM to accurately predict chromatin state at near single-cell resolution (related to Fig. 4). (a) Dot plot showing the percentage of non-empty bins across gene expression, accessibility, and 5mCG for seven major cell types at the pseudobulk, metacell, and single-cell levels. (b) Dot plot showing the percentage of non-empty bins across gene expression, accessibility, and 5mCG for seven major cell types across different genomic bin sizes (1 kb to 100 kb). (c) Six heatmaps showing emission probabilities for models with various numbers of inferred chromatin states, ranging from 3 to 8 states. (d) Heatmap showing enrichment of each chromatin state across genomic regions spanning upstream25 kb to downstream 25 kb around TSSs. (e) Heatmaps showing binarized input tracks from RNA expression, chromatin accessibility, and 5mCG across chromosome 1. Rows represent metacells. (f) Heatmap showing the relative genomic distances between pairs of inferred chromatin states. (g) Heatmap showing the distribution of chromatin states inferred from the full dataset. Annotations below the heatmap indicate CpG density, A/T content, lamina-associated domain (LAD) enrichment, and publicly reported EpiMap chromatin states.

**Extended Data Figure 5.**
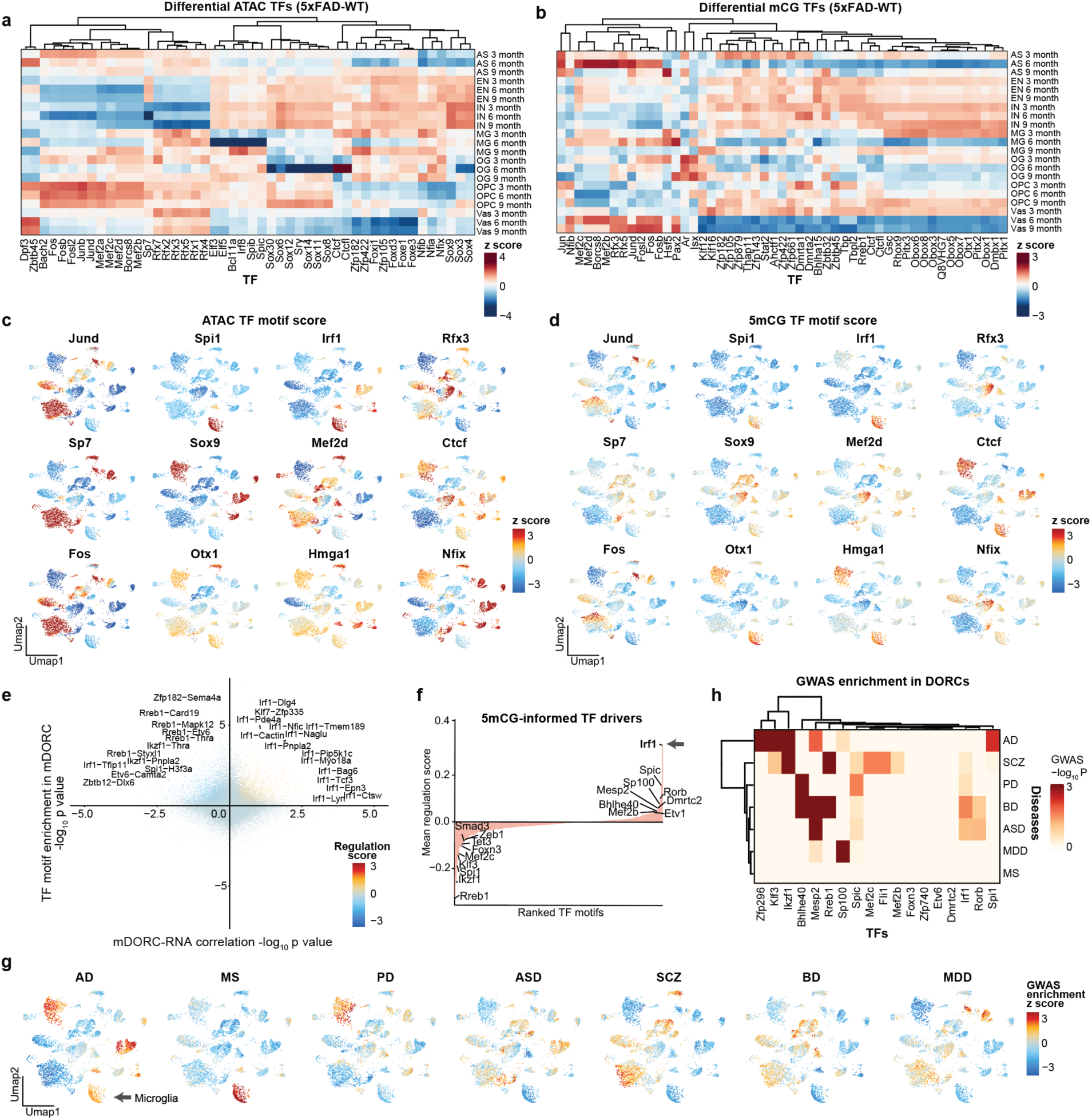
DNA methylation and accessibility provide complementary information inferring TF activity (related to Fig. 5). (a) Heatmap showing differences in TF motif scores inferred from chromatin accessibility between 5xFAD and WT mice. (b) Heatmap showing differences in TF motif scores inferred from 5mCG between 5xFAD and WT mice. (c,d) UMAP visualizations showing chromatin accessibility-inferred (c) and 5mCG-inferred (d) TF motif activity scores for 12 representative TFs across all profiled cell types. (e) Dot plot showing TF-mDORC associations, with the x-axis indicating the significance of correlation between TF expression and mDORC scores (*Z*-test relative to background) and the y-axis indicating relative enrichment of TF motifs among mDORC-associated peaks; points are colored by regulation score. (f) TFs ranked by their overall regulatory influence, quantified as the mean regulation score across all mDORCs. (g) UMAP visualizations showing the 5mCG inferred cell-type-specific enrichment of disease-associated GWAS variants. (h) Heatmap depicting the log-transformed enrichment P values of disease-associated GWAS variants within TF target gene–associated DORCs.

## Resource availability

### Materials availability

All unique/stable reagents generated in this study are available from the Lead Contact with a completed Materials Transfer Agreement.

## Acknowledgments

We thank all the members of the Ma labs for critical reading of the manuscript and helpful discussions. S.M acknowledges support from the NIH through awards U01CA290442, R61CA297881, and R21CA301237, and Mount Sinai FBI scholar. This work was supported in part by the research funding from NIH to K.L.C (R01AI168004).

## Author contributions

B.Z. and S.M. developed ME-seq protocol. B.Z. generated the single-cell sequencing library. A.G. collected mouse brain tissues. Z.L., M.Z, and E.W. performed sorting and sample processing for the fetal liver samples. A.K. and M.J.W. collected embryo samples. J.F. and P.R. provided human brain samples. F.D.T. provided other mouse tissues. M.A. performed functional validations of IRF1. B.Z., Z.W., J.L., and S.M. conducted the data analysis. A.G. and P.R. contributed to the interpretation of the data and the design of experiments. All authors participated in writing the manuscript. K.L.C., N.Y., and S.M. conceived and supervised the research, and all authors reviewed the manuscript.

## References

1 Lu, A. T. et al. Universal DNA methylation age across mammalian tissues. Nat Aging 3, 1144–1166 (2023). 10.1038/s43587-023-00462-6

2 Johnson, K. C. et al. Single-cell multimodal glioma analyses identify epigenetic regulators of cellular plasticity and environmental stress response. Nat Genet 53, 1456–1468 (2021). 10.1038/s41588-021-00926-8

3 Conole, E. L. S., Robertson, J. A., Smith, H. M., Cox, S. R. & Marioni, R. E. Epigenetic clocks and DNA methylation biomarkers of brain health and disease. Nat Rev Neurol 21, 411–421 (2025). 10.1038/s41582-025-01105-7

4 Chaligne, R. et al. Epigenetic encoding, heritability and plasticity of glioma transcriptional cell states. Nat Genet 53, 1469–1479 (2021). 10.1038/s41588-021-00927-7

5 Hou, Y. et al. Ageing as a risk factor for neurodegenerative disease. Nat Rev Neurol 15, 565–581 (2019). 10.1038/s41582-019-0244-7

6 Habib, N. et al. Disease-associated astrocytes in Alzheimer’s disease and aging. Nat Neurosci 23, 701–706 (2020). 10.1038/s41593-020-0624-8

7 Butovsky, O. & Weiner, H. L. Microglial signatures and their role in health and disease. Nat Rev Neurosci 19, 622–635 (2018). 10.1038/s41583-018-0057-5

8 Chen, S. et al. The global macroeconomic burden of Alzheimer’s disease and other dementias: estimates and projections for 152 countries or territories. Lancet Glob Health 12, e1534–e1543 (2024). 10.1016/S2214-109X(24)00264-X

9 Shireby, G. et al. DNA methylation signatures of Alzheimer’s disease neuropathology in the cortex are primarily driven by variation in non-neuronal cell-types. Nat Commun 13, 5620 (2022). 10.1038/s41467-022-33394-7

10 Lunnon, K. et al. Methylomic profiling implicates cortical deregulation of ANK1 in Alzheimer’s disease. Nat Neurosci 17, 1164–1170 (2014). 10.1038/nn.3782

11 De Jager, P. L. et al. Alzheimer’s disease: early alterations in brain DNA methylation at ANK1, BIN1, RHBDF2 and other loci. Nat Neurosci 17, 1156–1163 (2014). 10.1038/nn.3786

12 Corces, M. R. et al. Single-cell epigenomic analyses implicate candidate causal variants at inherited risk loci for Alzheimer’s and Parkinson’s diseases. Nat Genet 52, 1158–1168 (2020). 10.1038/s41588-020-00721-x

13 Xiong, X. et al. Epigenomic dissection of Alzheimer’s disease pinpoints causal variants and reveals epigenome erosion. Cell 186, 4422–4437 e4421 (2023). 10.1016/j.cell.2023.08.040

14 Morabito, S. et al. Single-nucleus chromatin accessibility and transcriptomic characterization of Alzheimer’s disease. Nat Genet 53, 1143–1155 (2021). 10.1038/s41588-021-00894-z

15 Mathys, H. et al. Single-cell multiregion dissection of Alzheimer’s disease. Nature 632, 858–868 (2024). 10.1038/s41586-024-07606-7

16 Liu, F., Wang, Y., Gu, H. & Wang, X. Technologies and applications of single-cell DNA methylation sequencing. Theranostics 13, 2439–2454 (2023). 10.7150/thno.82582

17 Smallwood, S. A. et al. Single-cell genome-wide bisulfite sequencing for assessing epigenetic heterogeneity. Nat Methods 11, 817–820 (2014). 10.1038/nmeth.3035

18 Luo, C. et al. Single nucleus multi-omics identifies human cortical cell regulatory genome diversity. Cell Genom 2 (2022). 10.1016/j.xgen.2022.100107

19 Itai, Y., Rappoport, N. & Shamir, R. Integration of gene expression and DNA methylation data across different experiments. Nucleic Acids Res 51, 7762–7776 (2023). 10.1093/nar/gkad566

20 Stuart, T. et al. Comprehensive Integration of Single-Cell Data. Cell 177, 1888–1902 e1821 (2019). 10.1016/j.cell.2019.05.031

21 Liu, Y. et al. Bisulfite-free direct detection of 5-methylcytosine and 5-hydroxymethylcytosine at base resolution. Nat Biotechnol 37, 424–429 (2019). 10.1038/s41587-019-0041-2

22 Ma, S. et al. Chromatin Potential Identified by Shared Single-Cell Profiling of RNA and Chromatin. Cell 183, 1103–1116 e1120 (2020). 10.1016/j.cell.2020.09.056

23 Olsen, T. R. et al. Scalable co-sequencing of RNA and DNA from individual nuclei. bioRxiv (2023). 10.1101/2023.02.09.527940

24 Jobson, D. D., Hase, Y., Clarkson, A. N. & Kalaria, R. N. The role of the medial prefrontal cortex in cognition, ageing and dementia. Brain Commun 3, fcab125 (2021). 10.1093/braincomms/fcab125

25 Network, B. I. C. C. A multimodal cell census and atlas of the mammalian primary motor cortex. Nature 598, 86–102 (2021). 10.1038/s41586-021-03950-0

26 Fumagalli, L. et al. Microglia heterogeneity, modeling and cell-state annotation in development and neurodegeneration. Nat Neurosci 28, 1381–1392 (2025). 10.1038/s41593-025-01931-4

27 Hou, J. et al. Transcriptomic atlas and interaction networks of brain cells in mouse CNS demyelination and remyelination. Cell Rep 42, 112293 (2023). 10.1016/j.celrep.2023.112293

28 Jin, K. et al. Brain-wide cell-type-specific transcriptomic signatures of healthy ageing in mice. Nature 638, 182–196 (2025). 10.1038/s41586-024-08350-8

29 Pishesha, N., Harmand, T. J. & Ploegh, H. L. A guide to antigen processing and presentation. Nat Rev Immunol 22, 751–764 (2022). 10.1038/s41577-022-00707-2

30 Persad, S. et al. SEACells infers transcriptional and epigenomic cellular states from single-cell genomics data. Nat Biotechnol 41, 1746–1757 (2023). 10.1038/s41587-023-01716-9

31 Chai, Y. L. et al. Lysosomal cathepsin D is upregulated in Alzheimer’s disease neocortex and may be a marker for neurofibrillary degeneration. Brain Pathol 29, 63–74 (2019). 10.1111/bpa.12631

32 Lancaster, T. M., Hill, M. J., Sims, R. & Williams, J. Microglia - mediated immunity partly contributes to the genetic association between Alzheimer’s disease and hippocampal volume. Brain Behav Immun 79, 267–273 (2019). 10.1016/j.bbi.2019.02.011

33 Deczkowska, A. et al. Disease-Associated Microglia: A Universal Immune Sensor of Neurodegeneration. Cell 173, 1073–1081 (2018). 10.1016/j.cell.2018.05.003

34 Ernst, J. & Kellis, M. ChromHMM: automating chromatin-state discovery and characterization. Nat Methods 9, 215–216 (2012). 10.1038/nmeth.1906

35 Boix, C. A., James, B. T., Park, Y. P., Meuleman, W. & Kellis, M. Regulatory genomic circuitry of human disease loci by integrative epigenomics. Nature 590, 300–307 (2021). 10.1038/s41586-020-03145-z

36 Auvray, C. et al. HOXC4 homeoprotein efficiently expands human hematopoietic stem cells and triggers similar molecular alterations as HOXB4. Haematologica 97, 168–178 (2012). 10.3324/haematol.2011.051235

37 Gao, T. et al. Transcriptional regulation of homeostatic and disease-associated-microglial genes by IRF1, LXRbeta, and CEBPalpha. Glia 67, 1958–1975 (2019). 10.1002/glia.23678

38 Johnston, R. A., Aracena, K. A., Barreiro, L. B., Lea, A. J. & Tung, J. DNA methylation-environment interactions in the human genome. Elife 12 (2024). 10.7554/eLife.89371

39 Yin, Y. et al. Impact of cytosine methylation on DNA binding specificities of human transcription factors. Science 356 (2017). 10.1126/science.aaj2239

40 El-Hodiri, H. M. et al. Nuclear Factor I in neurons, glia and during the formation of Muller glia-derived progenitor cells in avian, porcine and primate retinas. J Comp Neurol 530, 1213–1230 (2022). 10.1002/cne.25270

41 Nomaru, H. et al. Fosb gene products contribute to excitotoxic microglial activation by regulating the expression of complement C5a receptors in microglia. Glia 62, 1284–1298 (2014). 10.1002/glia.22680

42 Podlesny-Drabiniok, A. et al. BHLHE40/41 regulate microglia and peripheral macrophage responses associated with Alzheimer’s disease and other disorders of lipid-rich tissues. Nat Commun 15, 2058 (2024). 10.1038/s41467-024-46315-7

43 Novikova, G. et al. Integration of Alzheimer’s disease genetics and myeloid genomics identifies disease risk regulatory elements and genes. Nat Commun 12, 1610 (2021). 10.1038/s41467-021-21823-y

44 Hawrylycz, M. et al. SEA-AD is a multimodal cellular atlas and resource for Alzheimer’s disease. Nat Aging 4, 1331–1334 (2024). 10.1038/s43587-024-00719-8

45 Masuda, T. et al. Transcription factor IRF1 is responsible for IRF8-mediated IL-1β expression in reactive microglia. J Pharmacol Sci 128, 216–220 (2015). 10.1016/j.jphs.2015.08.002

